# Rescue of a developmental arrest caused by a *C. elegans* heat-shock transcription-factor mutation by loss of ribosomal S6-kinase activity

**DOI:** 10.1101/310086

**Authors:** Peter Chisnell, T. Richard Parenteau, Elizabeth Tank, Kaveh Ashrafi, Cynthia Kenyon

## Abstract

The widely conserved heat-shock response, regulated by heat shock transcription factors, is not only essential for cellular stress resistance and adult longevity, but also for proper development. However, the genetic mechanisms by which heat-shock transcription factors regulate development are not well understood. In *C. elegans*, we conducted an unbiased genetic screen to identify mutations that could ameliorate the developmental arrest phenotype of a heat-shock factor mutant. Here we show that loss of the conserved translational activator *rsks-1/S6-Kinase, a downstream effector of TOR kinase, can rescue the developmental-arrest phenotype of hsf-1* partial loss-of-function mutants. Unexpectedly, we show that the rescue is not likely caused by reduced translation, nor to activation of any of a variety of stress-protective genes and pathways. Our findings identify an as-yet unexplained regulatory relationship between the heat-shock transcription factor and the TOR pathway during *C. elegans’* development.

## Introduction

The health and longevity of an organism depends on robust proteostatic machinery to keep proteins functioning properly. One major source of cellular quality control is the heat-shock response. The heat-shock response increases expression of a variety of chaperones in response to many stresses, including heat and heavy metals. These so-called heat-shock proteins are coordinately regulated by one (in yeast and invertebrates) or multiple (in vertebrates) transcription factors called heat shock factors. In addition, heat shock factors act under normal conditions to promote longevity and developmental growth to adulthood.

In *S. cerevisiae, C. elegans*, and *Drosophila*, a loss of heat shock factor leads to developmental arrest at non-stressful temperatures. In yeast, the developmental arrest caused by lack of HSF-1 activity can be rescued by restoring basal expression levels of two heat shock proteins, Hsp70 and Hsp90, but additional Hsf1 target genes are required for resistance to heat stress (Solís *et al*. 2016). In *Drosophila*, loss of *hsf* causes arrest at the first or second larval-instar stage of development, as well as defects in oogenesis (Jedlicka *et al*. 1997). Unlike in yeast, Jedlicka *et al*. found that this essential development function in *Drosophila* was not mediated through canonical heat shock genes. In *C. elegans*, HSF-1 is needed for progression past the L2-L3 larval stage, and it has been shown to regulate a developmental program that is distinct from the heat shock response by binding to a promoter sequence different from the canonical sequence (Li *et al*. 2016).

Heat shock factors are important but not essential for mouse development. Mouse cells lacking Hsf1 or both Hsf1 and Hsf2 can be cultured and show normal constitutive transcription of heat shock proteins, although cells without Hsf1 are sensitive to heat stress (McMillan *et al*. 1998; Solís *et al*. 2016; Zhang *et al*. 2002). *In vivo*, Hsf1(-/-) mice display additional phenotypes, including growth retardation, female infertility, prenatal lethality (Xiao *et al*. 1999, Christians *et al*. 2000), and neurological defects (Santos and Saraiva 2004, Takaki *et al*. 2006). Hsf2(-/-) mice also display abnormalities, in both gametogenesis and brain structure (Chang *et al*. 2006).

While these studies establish that a heat-stress-independent activity of *hsf-1* is inextricably tied to development, the genetic pathways and mechanisms by which heat-shock factors promote growth and development remain largely unexplored. To address this question in an unbiased way, we conducted a genetic screen in *C. elegans* for suppressor mutations that allow *hsf-1(sy441)* mutant animals to grow to adulthood. The sy441-mutant HSF-1 protein lacks its transactivation domain, severely blunting its ability to regulate canonical heat-shock response genes (Hajdu-Cronin *et al*. 2004). These animals grow well at lower temperatures (15-20°C in our hands), but when grown at temperatures of 25°C or higher, they display a developmental phenotype similar to that of *hsf-1* loss-of-function mutants, arresting at the L2-L3 stage (Li et al. 2016).

In this study, we show that mutations in multiple complementation groups allow developing *sy441* mutants to reach adulthood. These mutations dramatically postpone, but do not eliminate, growth arrest within the strain’s lineage, as the rescued adults produce progeny that fail to progress through development. We show that one of these mutations is a putative null allele of rsks-1/S6Kinase, a translational activator regulated by TOR. We find that knockdown of various TOR-pathway components can also rescue the phenotype. Unexpectedly, we find no evidence that phenotypic rescue is mediated by inhibition of translation. Nor does rescue appear to be due to the activation of other stress responses such as the ER unfolded-protein response or the mitochondrial unfolded-protein response; nor is it due to the activation of various stress-resistance pathways known to extend adult lifespan.

We find that the rescue mediated by loss of S6 Kinase is dependent upon residual activity of the mutant HSF-1(sy441) protein, and that transgenically-increased expression of the *hsf-1(sy441)* allele is sufficient on its own to rescue development. However, we see no evidence that the S6 kinase mutation increases expression of *hsf-1* nor any of a variety of target genes we examined. We conclude that loss of S6 kinase could potentially elevate other targets of *hsf-1* or, alternatively, provide other factors that allow low levels of HSF-1 activity to sustain growth to adulthood.

## Materials and Methods

### C. elegans *strains*

Wild-type (N2)
VB654: *rsks-1(sv31)*; svEx136[*unc*-36(+) *rsks-1(+) sur-5::gfp*]
AGD794: *hsf-1(sy441); uthIs225*[*sur-5p::hsf-1(sy441); myo2p::tdTomato*]
CF3951: *hsf-1(sy441)*
CF4522: *hsf-1(sy441)* with an unidentified suppressor mutation
CF4523: *hsf-1(sy441)* with an unidentified suppressor mutation
CF4524: *hsf-1(sy441)* with an unidentified suppressor mutation
CF4525: *hsf-1(sy441); rsks-1(mu482)*
CF4526: *hsf-1(sy441)* with an unidentified suppressor mutation
CF4540: *hsf-1(sy441); rsks-1(sv31)*
CF4542: *hsf-1(sy441); rsks-1(mu482); svEx136[unc-36(+) rsks-1(+) sur-5::gfp]*
CF4543: *hsf-1(sy441); T24F1.4(tm5213)*

### Development assay

Arrested L1 larvae were spotted onto plates preheated to 25.8°C. After spotting, plates were placed back at 25.8°C in an open Tupperware container for one hour to re-equilibrate temperature. After one hour, the container was closed to prevent drying (except for one corner) for four days. Each condition utilized four plates each containing roughly 25 animals. After 4 days, conditions were blinded and scored on a qualitative five-point scale for developmental stage. Animals with a score of three had passed the point of vulval eversion (transition to adulthood) but had not yet produced any eggs. Therefore, animals with a score of three or higher were classified as adults in the figures and tables shown here.

### EMS Mutagenesis and screening

Mutant *hsf-1(sy441)* worms grown at 20°C on OP50 were bleach-prepped, and the eggs were incubated overnight in M9. The next day, a total of 10,000 L1 larvae were divided among four 10 cm plates containing OP50 (~2500/plate) and incubated at 20°C. Once this P_0_ population reached early L4, the worms were collected from the plates, washed 3 times, and re-suspended in 15 mL of M9 buffer. In a separate 50 mL conical tube, 100 uL of EMS (Sigma #M-0880) was mixed into 5 mL of M9. The worm suspension was then added to this new conical tube containing EMS, the top was parafilmed, and the tube was placed in a rotator at 20°C for 4 hours. The final concentration of EMS under these conditions was ~50 mM, for an expected mutation rate of 5x10^-4^ mutations/gene/gamete (Brenner, 1974). Next, the worms were pelleted, the supernatant was removed, and the worms were washed thrice with 15 mL of M9. Lastly, the worm pellet was split equally among four 10 cm plates containing OP50, and recovered at 20°C for 24 hours.

At the end of this recovery period, the P_0_ animals were collected from the plates and bleached using standard protocol. Assuming 5 viable eggs were obtained per P_0_, this yielded a total of ~50,000 FI’s, representing 50x genomic coverage. These eggs were plated evenly onto 20 separate 10 cm plates containing OP50 (2500/plate), and grown to adulthood at 20°C. The gravid populations on each of these 20 plates were then collected individually and bleached, and each resulting batch of F_2_ eggs was plated onto its own 15 cm plate containing OP50 (25,000/plate). The F_2_ worms were then grown at 25°C. On day 5, the plates were screened for any worms that had developed beyond the L3 arrest phase, and these candidates were collected onto individual 3cm OP50 plates and recovered at 20°C.

### Validation and Phenotypic Analysis of Screen Hits

All candidates recovered from the screen were validated by determining whether the suppression-of-hsf-1-arrest phenotype bred true to the next generation. Those that failed this test were discarded. The validated lines were then genotyped for reversal of the *hsf-1(sy441)* point mutation, with intent to discard true genetic revertants.

In order to narrow down the 17 remaining mutant lines, we further characterized them for penetrance and expressivity. First, we measured the percentage of suppression-of-hsf-1-arrest that each line displayed. This was done by picking 50 eggs onto a 3 cm OP50 plate, incubating for 4 days at 25°C, and then counting the number of worms that developed past L3 arrest. Investigation was continued only for lines with 50% or higher population rescue. Next, we determined the furthest developmental stage each line could achieve at 25°C, using the same experimental set-up just described, and only pursued those that could produce gravid adults.

### Genetic Characterization of Screen Hits

We characterized the allelic nature of the 9 remaining lines. Males were generated for each, as well as for *hsf-1(sy441)* single mutants, and reciprocal crosses were carried out to identify any dominant or sex-specific mutations. Given that all suppressors were found to be recessive, the males generated above were then used to perform complementation assays. To ensure independence, all lines isolated from the same plate (of the original 20) were interrogated with reciprocal crosses, looking for heterozygous progeny that maintained the suppression-of-hsf-1-arrest phenotype. Second, reciprocal complementation crosses were carried out between all remaining lines (across the original 20 populations), with the intent to again keep only one from each complementation group. This yielded a set of five lines with unique, recessive, and penetrant mutations that allowed *hsf-1(sy441)* worms to reach gravid adulthood at 25°C. Four of these five strains exhibited extended lifespan at 20°C, but this phenotype was lost upon backcrossing.

### Identification of candidate genes

These five suppressor strains were backcrossed to the control *hsf-1(sy441)* strain a total of six times before genomic sequencing. The suppressor strains were then backcrossed a total of nine times before use in all other experiments.

Genomic DNA from each mutant strain was isolated from roughly 300 ul of pelleted worms from a mixed population. DNA was sheared using the Covaris M220 sonicator to an average length of 350 base pairs. After shearing, DNA was quantified using the Qubit dsDNA high sensitivity assay kit. DNA libraries were prepared from genomic DNA utilizing the Bioo Scientific NEXTflex Rapid DNA-Seq Library Prep Kit. Library quality was assessed utilizing the Agilent high-sensitivity DNA analysis kit. Libraries were sequenced by the UCSF Institute for Human Genetics Core Facility according to the manufacturer’s protocol using an Illumina HiSeq 2500. Reads were analyzed using Amazon Web Services.

Genomic information from the Human Genetics Core Facility was analyzed using the Galaxy platform. Genes with high-quality protein-coding point mutations were then examined for effects on the development arrest phenotype.

### Lifespan Analysis

Life span analysis was performed at 20°C as described previously (Apfeld and Kenyon 1998). Animals were grown on OP5O *E. coli* and transferred to fresh plates on the first day of adulthood (“day 1”) and every 2 days thereafter. Every two days, starting on day 1, animals were scored. Animals that moved were scored as alive, animals that did not move, even after being prodded with a platinum wire, were scored as dead, and animals that could not be found or displayed phenotypes such rupturing or bagging were censored.

### Translation Inhibiting Drugs

Salubrinal (SML0951) and homoharringtonine (SML1091) were purchased from Sigma-Aldrich. Cycloheximide (#94271) was purchased from VWR Life Sciences. Drugs were diluted and spotted on plates with bacteria to a final DMSO concentration of 0.19% (unless otherwise noted) and allowed to diffuse into the agar for one day before animals were plated for development experiments.

### Real-time qPCR

RNA extraction, cDNA preparation, and real-time qPCR were performed as described previously (Van Gilst *et al*. 2005) from 3 biological replicates of each condition, with at least 1,000 animals included in each condition. Animals were grown from arrested L1 larvae at 25.8°C until >50% of the population had grown to the lethargus stage between the L1 and L2 larval stages (approximately 13 hours for wild-type background and approximately 16 hours for *hsf-1(sy441))* background). Data were standardized to three control primers: *tba-1,cdc-42*, and *pmp-3*. Primer sequences are listed in table Supplementary Table 3.

### RNAi

HT-115 RNAi bacteria were obtained from existing Ahringer and Vidal libraries. The control for all RNAi experiments was L4440 vector. For each experiment, single colonies were grown overnight at 37°C in 3 mL LB with 100μg/mL carbenicillin and 10μg/mL tetracycline. The next day, the resulting stationary-phase cultures were diluted tenfold with LB containing 100μg/mL carbenicillin and grown for two hours at 37°C. 1M IPTG was then added to the culture to reach a final concentration of 2mM, after which 100 μL of the resulting culture were spotted onto NG plates. The bacteria were grown on these plates overnight at 30°C. Animals were placed on plates the following day in all experiments except those involving a drug condition, in which case drugs were added and left to diffuse into the plate for an additional day before the addition of experimental and control animals.

### Statistical Analyses

On all bar graphs, bars represent the mean, and error bars represent the SEM. For developmental assays of multiple genes in parallel, significance was measured across three independent experiments using Cochran-Mantel-Haenszel tests with Bonferroni corrections unless otherwise noted. For lifespan assays, log-rank tests with Bonferroni corrections were used. For RT-qPCR experiments, one-way ANOVAs with Tukey post tests were used on three biological replicates.

### Data Availability

Strains are available upon request. The authors state that all data necessary for confirming the conclusions presented in the article are represented fully within the article.

## Results

### A genetic screen identifies mutations that rescue the hsf-1(sy441) developmental arrest

To investigate how *hsf-1* regulates development, we conducted an unbiased forward mutagenesis screen in *C. elegans* to rescue the growth-arrest phenotype displayed by *hsf-1(sy441)* mutants grown at 25°C. Following EMS mutagenesis (Brenner 1973), we recovered seventeen independent mutants that, when shifted from 20°C to 25.8°C at the mid-L4 (final) larval stage, produced progeny that grew to adulthood. We picked five strains (see methods) to backcross and sequence to discover candidate genes. *hsf-1(sy441)* mutants carrying the suppressors reached adulthood exhibiting apparently-normal anatomy, but they displayed a reduced body size and could not be cultivated past a single generation at 25.8°C (Figure 1A, B). The progeny of some strains grown at 25.8°C produced eggs that did not hatch, whereas others produced progeny that arrested at L1. If adult animals were shifted back to 20°C, they produced offspring which developed normally.

**Figure 1:**
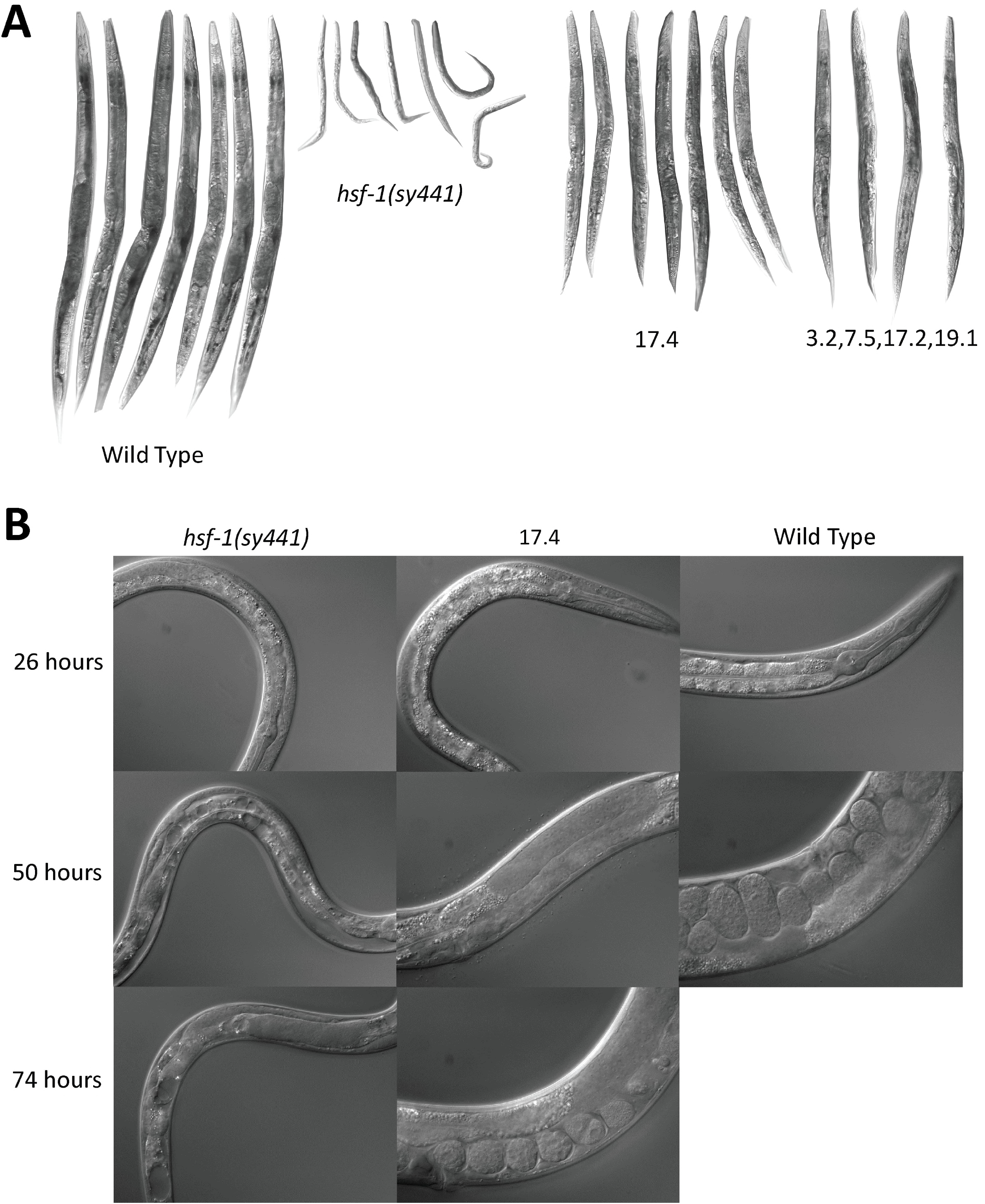
Mutants discovered in a mutagenesis screen for rescue of the developmental arrest of *hsf-1(sy441)*. (A) Appearance of wild-type, *hsf-1(sy441)*, and multiple suppressor strains after 4 days at 25.8°C from arrested L1 larvae extracted at 20°C, imaged at 100x magnification. Strains CF4522, CF4523, CF4524 and CF4526 contain additional suppressors from the screen, but were not analyzed further in this study. (B) Appearance of wild-type, *hsf-1(sy441)*, and CF4525 suppressor strain grown at 25.8°C to various time-points from arrested L1 larvae extracted at 20°C, imaged at 600x magnification.

When parents were shifted to 25.8°C at the L4 stage, 100% of *hsf-1(sy441)* progeny without suppressor mutations arrested at the L1-L3 stage, whereas 100% of progeny homozygous for the five suppressor mutations reached adulthood. If instead eggs were extracted from the gonads of the parental generation at 20°C, allowed to hatch into minimal medium (causing L1 developmental arrest), and then placed on 25.8°C plates, a small percentage of *hsf-1(sy441)* single mutant-animals could reach early adulthood but rarely produced eggs, while still 100% of progeny homozygous for the suppressor mutations reached adulthood. In order to detect subtle shifts in the percentage of animals that reached adulthood, all of the assays described here utilized arrested L1 larvae hatched at 20°C and then shifted to higher temperature.

Because our suppressor mutations rescued the developmental growth-arrest phenotype of *hsf-1(sy441)* animals, we tested whether they could also rescue another well-documented *hsf-1(sy441)* phenotype, shortened adult lifespan (Garigan *et al*. 2002; Hajdu-Cronint et al 2004). None of the mutants extended the shortened lifespan that *hsf-1(sy441)* animals exhibit at the temperatures permissive for growth (Supplementary Figure 1).

### *Loss of the gene* rsks-1 *rescues the* hsf-1(sy441) *development arrest*

Genomic sequencing revealed that one of the suppressor strains (CF4525) contained a premature stop codon in the gene *rsks-1* at the 295^th^ base pair, *rsks-1(mu482)*. The *rsks-1* gene encodes *C. elegans*’ ribosomal S6 Kinase ortholog, which promotes normal levels of translation. We determined that this mutation was the causative factor for the developmental rescue in two ways: first, we confirmed the developmental rescue with a previously-isolated null allele, *rsks-1(sv31)*, and with RNAi-mediated knockdown of *rsks-1* (Figure 2A). Second, we showed that transgenic re-expression of *rsks-1* in the suppressor strain blocked the rescue phenotype (Figure 2B, C).

**Figure 2:**
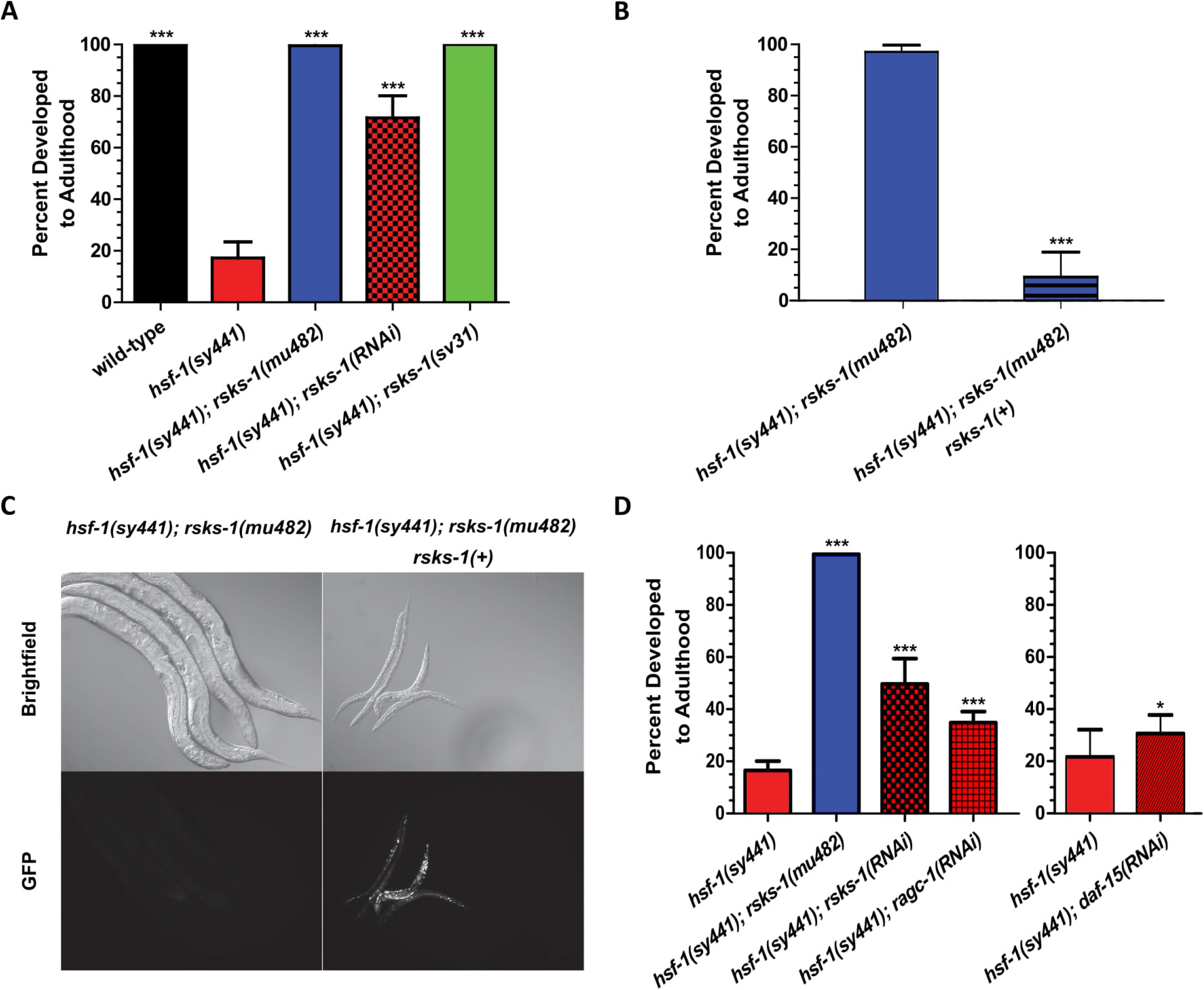
Loss of *rsks-1/S6K* or reduction of TOR function was sufficient to rescue the developmental arrest of *hsf-1(sy441)*. (A) Loss of *rsks-1* rescued the developmental arrest of *hsf-1(sy441)*, ***p < 0.001 compared to *hsf-1(sy441)* using the CMH test with three experiments. (B,C) Transgenic overexpression of *rsks-1::GFP* in *hsf-1(sy441); rsks-1(mu482)* double mutants suppressed the rescue. All animals are *hsf-1(sy441); rsks-1(mu482)* double mutants with the stochastically-inherited transgene *svEx136[unc[36(+) rsks-1(+) sur-5::gfp]*. Strain identity was blinded and development was scored, and then groups were categorized based on whether GFP was visible, ***p<0.001 compared to animals without GFP using the CMH test with three experiments; animals were imaged at 100x magnification. (D) RNAi knockdown of genes in the TOR pathway also rescued the developmental arrest of *hsf-1(sy441)*, *p<0.05, ***P<0.001 compared to *hsf-1(sy441)* using the CMH test with three experiments (four for *daf-15)*.

### *Reduction in TOR-pathway function also rescues* hsf-1(sy441) *development*

What is the mechanism by which *rsks-1* loss influences *hsf-1* mutants? S6 kinase is a key target of TOR kinase. To determine if the developmental rescue phenotype could be produced by a reduction in TOR activity more broadly, we used RNAi to knock down daf-15/RAPTOR, a component of the TORC1 complex, as well as *ragc-1/RAG* GTPase, a positive regulator of TORC1 (Fukuyama *et al*. 2012; Jia *et al*. 2004). Reduced expression of both of these genes significantly rescued the *hsf-1(sy441)* developmental arrest (Figure 2D), albeit to a lower degree than did RNAi knockdown of *rsks-1*.

### rsks-1(-)-*mediated* hsf-1(sy441) *rescue was not mimicked by a reduction in translation*

Loss of *rsks-1* causes a reduction in translation (Hansen *et al*. 2006). One possible explanation for the mechanism of developmental rescue conferred by *rsks-1* mutation is that *hsf-1(sy441)* animals, predicted to have reduced proteostatic capability, are overburdened by the amount of protein being translated during development at 25°C, and the reduced rate of translation incurred by mutating *rsks-1* reduces that burden to a manageable level in *hsf-1(sy441)* worms. To investigate this hypothesis, we reduced translation in other ways, by treatment with several chemical inhibitors of translation and via various RNAi treatments. If our hypothesis were correct, one would expect the following: at high concentrations, these drugs would be lethal and, at low concentrations, they would have no effect, but at some range of intermediate concentrations, they would lower the translation rate to a level permissible for developmental rescue. The chemical inhibitors tested were salubrinal, which blocks translation initiation, and harringtonine and cycloheximide, which block translation elongation. For each concentration of chemical inhibitor, we quantified the percentage of animals that reached adulthood (Figure 3A-D, Supplementary Table 1). We also used RNAi to reduce translation, diluting RNAi with vector control to achieve multiple levels of knockdown. We RNAi-inhibited *ifg-1*, which promotes translation additively with S6 kinase (Pan *et al*. 2007), and *rps-6*, the small ribosomal protein that is phosphorylated by S6 kinase. Only *hsf-1(sy441)* animals grown on 46.9nM salubrinal showed a marginally-significant amount of development to adulthood (two other conditions, salubrinal at 375nM and *ifg-1* RNAi bacteria diluted 1:50 with vector-only control bacteria, neared significance with p values of 0.055 and 0.088, respectively). None of these conditions produced an effect similar to that of the *rsks-1* null mutation in terms of either penetrance or expressivity, leading us to believe that reduced translation plays little to no role in relieving the block in *hsf-1(sy441)’s* development.

**Figure 3:**
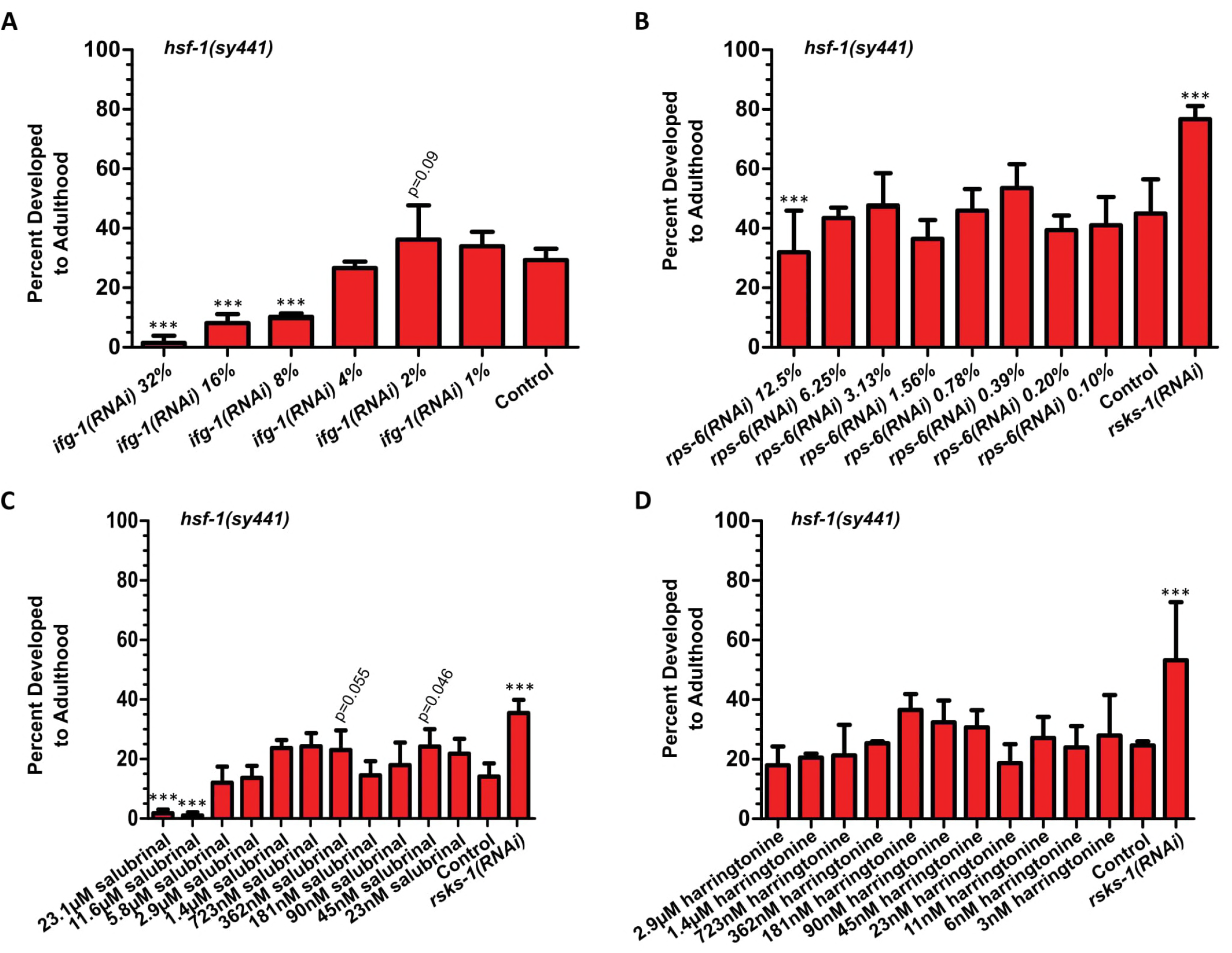
The ability of *rsks-1(-)* to rescue of the development of *hsf-1(sy441)* mutants is not explained by a reduction in translation. (A) None of multiple dilutions of *ifg-1* RNAi-bacteria with vector-control RNAi-bacteria rescued the developmental arrest (1% denotes 1:99 *ifg-1*:vector-control RNAi-bacteria), ***p<0.001 compared to control *hsf-1(sy441)* by CMH test with three replicates. (B) None of multiple dilutions of *rps-6* RNAi-bacteria with vector-control RNAi-bacteria rescued developmental arrest (1% denotes 1:99 rps-6:vector control RNAi), ***p<0.001 compared to control *hsf-1(sy441)* by CMH test with three replicates. (C) Two concentrations of the translation initiation inhibitor salubrinal showed a small amount of rescue near p=0.05 level of significance, ***p<0.001 compared to control *hsf-1(sy441)* by CMH test with three replicates. (D) None of multiple concentrations of the translation elongation inhibitor harringtonine rescued developmental arrest, ***p<0.001 compared to control *hsf-1(sy441)* by CMH test with two replicates.

### rsks-*1(-) mutants do not activate canonical unfolded-protein response pathways*

Because *rsks-1* mutation does not seem to rescue the *hsf-1(sy4441)* developmental arrest by lowering the translational burden on the animal, we hypothesized that it might be *increasing* the animal’s capability to manage proteotoxic stress through non-heat-shock response pathways such as the mitochondrial or ER unfolded-protein response (UPR) pathways. Like the heat-shock response, these pathways upregulate the expression of protein chaperones such as BIP/HSP-70 (Shen *et al*. 2001).

The development of *hsf-1(sy441); rsks-1* double mutants was partially inhibited by knockdown of the ER-UPR pathway gene *xbp-1* (Figure 4A), an intervention that has no effect on wild type, showing that the animals are sensitive to disruption of additional proteostatic systems beyond the heat-shock response. This suggests the possibility that *rsks-1* loss up-regulates *xbp-1* transcriptional-target genes, such as *hsp-4* (the *C. elegans* BIP ortholog). However, when assayed using RT-qPCR, *hsp-4* did not appear to be upregulated by loss of *rsks-1* in *hsf-1(sy441)* mutants (Figure 4B). Likewise, RT-qPCR of the mitochondrial-UPR target gene *hsp-6* revealed that the mitochondrial-UPR was not upregulated. These findings suggest that an increase in UPR function is not the mechanism by which *rsks-1* rescues development (Figure 4B).

**Figure 4:**
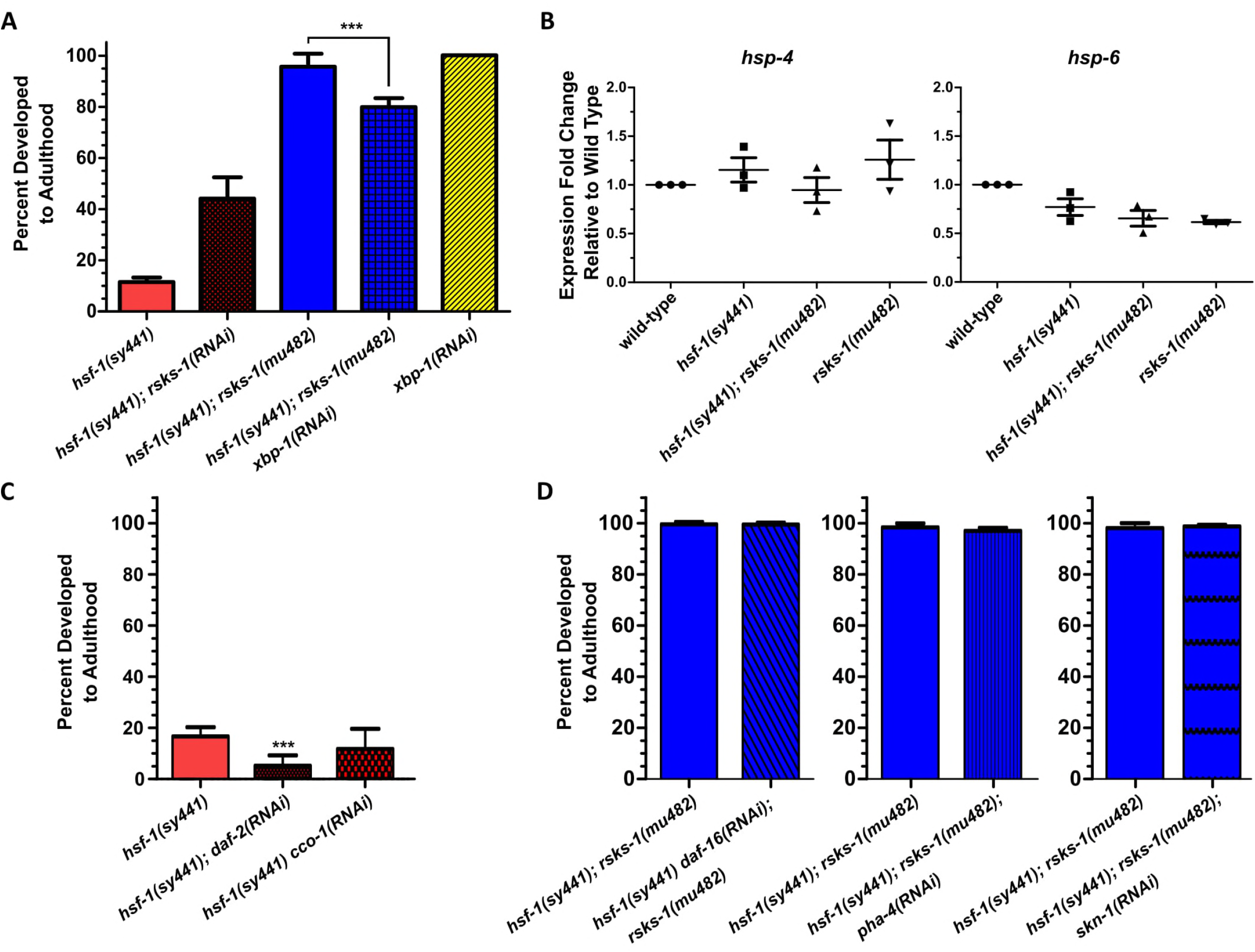
Stress-responses in *hsf-1(sy441)* animals carrying the *rsks-1(mu482)* suppressor. (A) *hsf-1(sy441); rsks-1(mu482)* double mutants are more sensitive to RNAi of *xbp-1*, an activator of the ER UPR, than are wild-type animals, ***p<0.001using CMH test with three trials. Animals were grown on HT115 RNAi or control empty-vector RNAi bacteria. (B) The canonical ER UPR and mitochondrial UPR chaperones were not upregulated in *hsf-1(sy441); rsks-1(mu482)* double mutants. (C) Inhibition of the *daf-2* insulin/IGF-1 receptor or the mitochondrial electron transport chain, which extend the lifespan of wild type, failed to rescue the developmental arrest of *hsf-1(sy441)* mutants. (D) Transcription factors required for TOR reduction-of-function to extend lifespan are not required for *rsks-1* mutation to rescue *hsf-1(sy441)* developmental arrest, using the CMH test with three trials.

### *The developmental rescue of* hsf-1(sy441) *mutants is TOR-specific but does not act through canonical lifespan pathways that interact with TOR*

TOR and heat shock factor are intimately tied to lifespan regulation and stress resistance. Because translation and UPR pathways did not seem to be the mechanisms through which *rsks-1* mutation rescued *hsf-1(sy441)’s* development, we wondered whether activating other pathways known to increase lifespan and proteostasis could also rescue its development. Both *daf-2* (insulin/IGF-1 receptor) and *cco-1* (mitochondrial electron transport gene) RNAi extend lifespan (Dillin *et al*. 2002; Kenyon *et al*. 1993), but neither rescued the block in *hsf-1(sy441)* development (Figure 4C). In fact, *daf-2* RNAi seemed to have a negative effect on development. To further investigate, we used RNAi to inhibit the transcription factors *daf-16, pha-4* and *skn-1*. Each of these genes promotes *C. elegans’* stress resistance and lifespan extension in TOR reduction-of-function and/or other long-lived mutants (Robida-Stubbs *et al*. 2012; Seo *et al*. 2013; Sheaffer et al. 2008), but none blocked the ability of *rsks-1* loss to rescue the *hsf-1(sy441)* developmental arrest (Figure 4D). We then tested a subset of genes in the published literature that have been shown to be a part of the genetic networks associated with *rsks-1* or *hsf-1*, including those from microarray datasets (Baird *et al*. 2014; Chen *et al*. 2013; Magnuson *et al*. 2012). Of the 75 genes tested, we saw no evidence of RNAi knockdown either rescuing the development of *hsf-1(sy441)* mutants or having a more severe reduction-of-development effect on *hsf-1(sy441); rsks-1* double mutants vs. wild-type (Supplementary Table 2).

### Developmental rescue requires residual hsf-1 function but does not act through canonical heat shock-regulated genes

Despite the lack of its transactivation domain, the fact that the *hsf-1(sy441)* allele does not cause arrest at 20°C, as the null mutation does (Li *et al*. 2016), shows that the HSF-1(sy441) mutant protein retains some functionality. To test whether endogenous levels of the mutant protein are required for *rsks-1* null mutations to rescue development at 25.8°C, we knocked down *hsf-1* using RNAi in the *hsf-1(sy441); rsks-1(mu482)* double mutant. RNAi knockdown of residual HSF-1(sy441) function blocked the *rsks-1* rescue phenotype (Figure 5A). In addition, we found that transgenically increasing the *hsf-1(sy441)* gene dosage was sufficient to rescue the developmental arrest of *hsf-1(sy441)* mutants (Figure 5B). However, despite the fact that residual levels of HSF-1(sy441) are necessary for developmental rescue and overexpression of HSF-1(sy441) can be sufficient for developmental rescue, we saw no evidence through RT-qPCR that *rsks-1* mutation elevated the expression levels of the *hsf-1* gene nor genes known to be regulated by HSF-1 in heat shock or developmental contexts (Figure 5C) (Li *et al*. 2016). Thus, the hypothesis that, directly or indirectly, loss of S6 kinase promotes growth to adulthood by enhancing the effectiveness of HSF-1(sy441) remains an attractive, though unproven, model.

**Figure 5:**
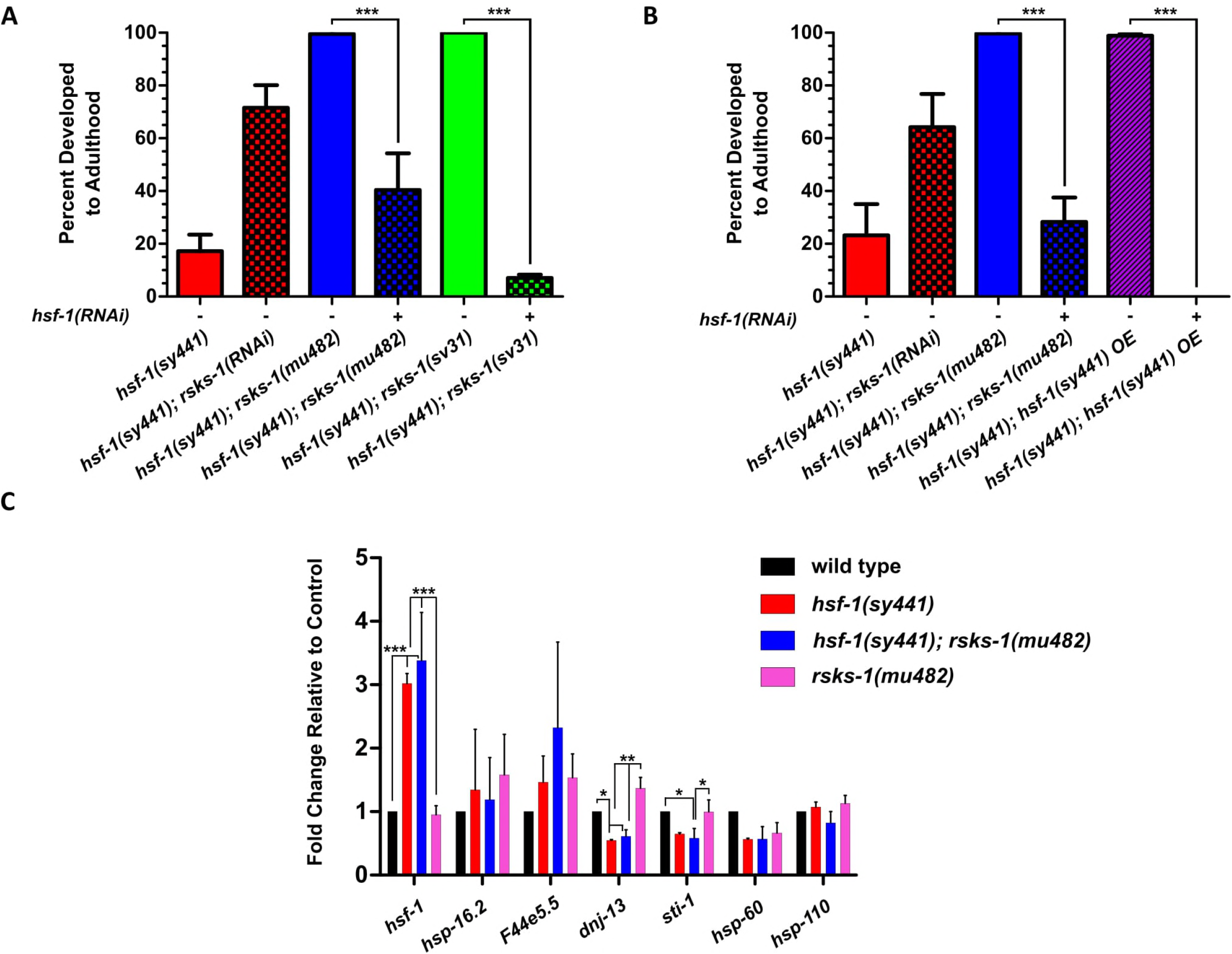
*hsf-1(sy441)* activity is necessary and can be sufficient to rescue developmental arrest, but *rsks-1(mu482)* does not appear to affect expression of *hsf-1* or its canonical targets. (A) *hsf-1* RNAi treatment prevented two *rsks-1/S6K* null mutations from rescuing the developmental arrest of *hsf-1(sy441)* mutants, ***p<0.001 using the CMH test with three trials. (B) Overexpression of the *hsf-1(sy441)* allele rescued the developmental arrest of *hsf-1(sy441)* mutants, ***p<0.001 using the CMH test with four trials. (C) Expression levels of canonical heat-shock genes and developmentally-regulated hsf-1-response genes are unaffected by the *rsks-1(mu482)* mutation, as measured by RT-qPCR, *p<0.05**p<0.01 ***p<0.001 by one-way ANOVA with Tukey multiple comparison post-test measured for each gene independently with three biological replicates.

## Discussion

*hsf-1* and its orthologs have been studied extensively in the contexts of stress, aging, and human pathologies such as cancer and neurodegenerative diseases (Li *et al*. 2017). Despite characterization of the requirement for a functioning heat shock factor during development, the specific molecular roles it plays in development remain largely unknown. In this paper we show that the TOR pathway, and more specifically the downstream TOR target ribosomal S6 kinase, acts to truncate development in *hsf-1(sy441)* reduction-of-function mutants.

Interestingly, lifespan extension caused by inhibiting S6-kinase or TOR activity is known to be blocked by the *hsf-1(sy441)* mutation (Seo *et al*. 2013). Consistent with this finding, our S6-kinase suppressor mutation failed to extend the lifespan of *hsf-1(sy441)* mutants. While that finding shows that HSF-1 is required for *rsks-1* loss to extend lifespan, here we find another reciprocal relationship; namely, that *rsks-1* loss can rescue the developmental arrest caused by reduction of HSF-1 function. In one case, low levels of S6K cause endogenous levels of HSF-1 to extend lifespan, whereas in the other case, low levels of HSF-1 cause endogenous levels of S6K to arrest development. The details of these reciprocal interactions remain elusive; nevertheless, our findings reveal a recurring, but potentially complex, relationship between these two important regulators of development, stress resistance, and lifespan.

It is interesting to note that when *hsf-1(sy441)* homozygotes are shifted to 25°C at the L4 larval stage, they lay eggs that hatch but fail to reach adulthood. However, when L1 larvae are starvation-arrested at 20°C and then fed and shifted to 25.8°C, some of those larvae reach adulthood. In other words, the general “arrest point” in the life cycle is different under these two conditions. This finding indicates that *hsf-1* loss does not cause a growth blockade at one specific point in the life cycle. Instead, this finding suggests the model that loss of HSF-1 function impacts a time-dependent accumulation of proteostatic damage rather than a specific developmental requirement. The opposite situation occurs in *Drosophila*, where HSF-1 activity is dispensable specifically after passing the first two larval stages (Jedlicka *et al*. 1997). Because no *hsf-1(sy441); rsks-1(mu482)* double-mutant animals arrest in the first generation but do produce progeny which then arrest, we hypothesize that loss of *rsks-1* does not bypass HSF-1 function but rather delays or reduces the damage caused by the reduction of HSF-1 function.

It seemed surprising that we were unable to phenocopy the effects of *rsks-1* mutation through any mechanism other than TOR knockdown, a condition predicted to reduce *rsks-1* activity. While we cannot know for certain whether a very specific amount of translation inhibition could rescue the developmental arrest, we have established that multiple methods of reducing translation at multiple concentrations are insufficient to rescue development. If reduced translation were the mechanism of growth arrest suppression, then the situation we found in this study would contrast dramatically with the effects that a similar scan of translation-RNAi knockdowns produced on another phenotype, lifespan extension. In that case, a wide variety of RNAi knockdowns scored positively, even without careful dose-response analysis (Hansen *et al*. 2006).

Another potential way that translation could be involved in this phenotype is if inhibiting S6K fails to inhibit, or even promotes, the translation of a specific subset of genes. Many stress responses, such as the heat shock response and integrated stress response, activate a subset of genes while inhibiting translation generally. If S6K inhibition causes a similar effect, then chemical or RNAi-mediated inhibition of translation alone, without activation of specific targets, would not be enough to rescue development.

Other than the small inhibition of rescue caused by *xbp-1* knockdown and the rescue caused by knockdown of *daf-15* and *ragc-1*, we did not identify other genes known to interact with *rsks-1* that are related to this phenotype. The finding that *daf-2* and *cco-1* knockdown do not rescue development makes it unlikely that the rescue is mediated by some general increase in stress resistance or lifespan-increasing pathways. Even so, it was surprising to us that knockdown of transcription factors *daf-16, pha-4, skn-1* failed to prevent the rescue phenotype. These transcription factors are essential for stress resistance and lifespan extension produced by inhibiting components of the TOR pathway. In addition, we saw no evidence across 85 genes, either predicted to be targets of S6K or proteins known to interact functionally with S6K, of RNAi inhibition rescuing development of *hsf-1(sy441)* mutants, or having a more severe effect on the development of *hsf-1(sy441); rsks-1* double mutants vs. wild-type. While we cannot be sure from these experiments of the efficacy of each individual RNAi clone, in each experiment positive controls were present, and in some cases a visible phenotype (other than rescue) was produced by the RNAi clone, such as loss of eggs in animals with inhibited *pha-4* (Supplementary Table 2).

Despite *rsks-1’s* requiring some level of *hsf-1* function to rescue the *hsf-1(sy441)* arrest phenotype, we saw no evidence of an activation of known hsf-1-activated genes or other stress-response pathways. In yeast, only two heat-shock proteins are required to rescue the loss of *hsf-1* for development (Solís *et al*. 2016), therefore it is possible that one or more unmeasured or non-canonical heat shock proteins are independently regulated by *hsf-1* and *rsks-1*, and it is through this unknown gene(s) that development is being rescued.

Most of the experiments in this study involved either modulating the expression levels of genes or measuring expression levels through RT-qPCR. As a kinase, it is possible that protein S6K modulates the activity of proteins directly without affecting expression levels. If such an activity increase activated the heat shock response, the ER UPR, or the mito-UPR broadly, we would expect to measure that through an increased expression of canonical target genes, but it is possible that S6 kinase phosphorylates specific proteins in these pathways to mediate developmental arrest. This would be interesting to investigate in the future.

In conclusion, a genetic screen for suppressors of the developmental growth arrest of *hsf-1(sy441)* partial loss-of-function mutants revealed a previously unknown relationship between heat-shock factor and TOR/S6 kinase activity. In some way, reducing S6 kinase activity allows animals with insufficient HSF-1 activity to progress much further through the life cycle than would otherwise be possible. Because heat-shock factor is known to enhance proteostasis under conditions of heat stress, we propose that loss of S6 kinase postpones arrest either by reducing the levels of damaged macromolecules produced in the cell, or by increasing the cell’s ability to remove or repair them.

---

**Supplementary Table 1:** Table of conditions that reduced translation.

**Supplementary Table 2:** RNAi knockdown of many genes associated with TOR, *rsks-1*, and *hsf-1* failed to rescue the *hsf-1(sy441)* developmental arrest or failed to prevent the *rsks-1(mu482)* mutation from rescuing the *hsf-1(sy441)* developmental arrest.

**Supplementary Table 3:** RT-qPCR primers

**Supplementary Figure 1:**
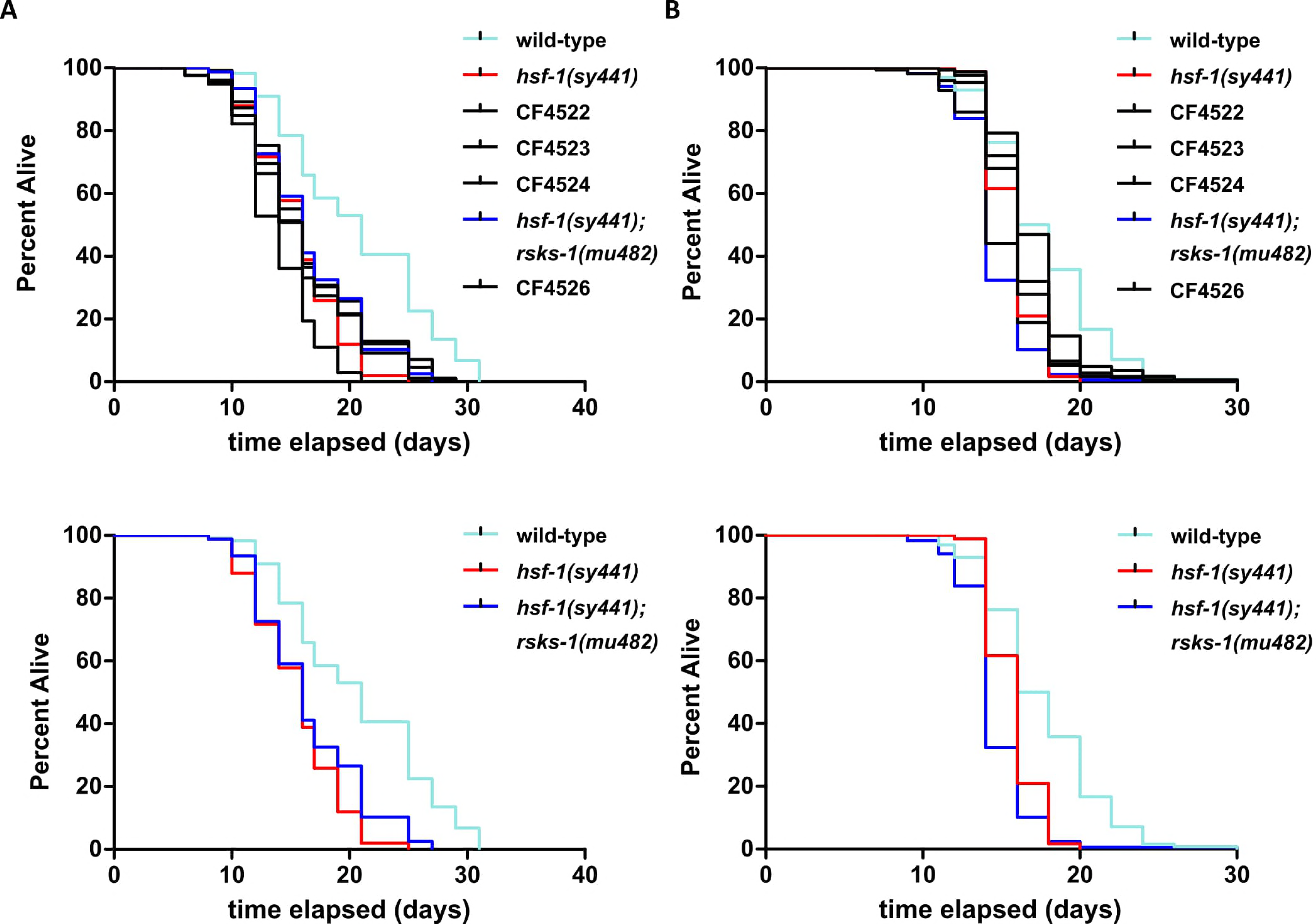
Strains carrying suppressors of the developmental arrest of *hsf-1(sy441)* mutants do not extend lifespan, either without (A) or with (B) the addition of 50μM FUDR to prevent progeny production. Bottom graphs display the same data as the graphs above them, showing only a subset of strains for clarity.

## Acknowledgements

The authors thank the *Caenorhabditis* Genetics Center, which is funded by the National Institutes of Health (NIH) Office of Research Infrastructure Programs (P40 OD-010440) for strains provided, Shohei Mitani of the National Bioresource Project for providing us with the *T24F1.4(tm5213)* allele, Simon Tuck of the Umea Center for Molecular Medicine for for providing us with the strain VB654, and Wormbase. The work was supported by NIH Ro1 grants AG011816 and AG046400.

